# Starting antiretroviral therapy within seven days of a positive HIV test increased the risk of loss to follow up in a primary healthcare clinic: a retrospective cohort study in Masaka, Uganda

**DOI:** 10.1101/640516

**Authors:** Julius Kiwanuka, Jacinta Mukulu Waila, Kahungu Methuselah Muhindo, Jonathan Kitonsa, Noah Kiwanuka

## Abstract

**Background:** Retention of patients initiated on antiretroviral therapy (ART) and good adherence remain cornerstones to long-term viral suppression. In this era of test and treat (T&T), ensuring that patients initiated on ART remain connected to HIV clinics will be key to the achievement of the UNAIDS 90-90-90 targets. Currently, limited studies have evaluated the effect instant ART initiation has on loss to follow up in a typical service healthcare setting. We studied the cumulative incidence, incidence rate of loss to follow up (LTFU), and factors associated with loss to follow up (LTFU) in a primary healthcare clinic that has practiced test and treat since 2012.

**Methods:** We retrospectively drew routine program data of patients initiated on ART from January 2012 to December 2016. We defined LTFU as failure of a patient to return to the HIV clinic for at least 90 days from the date of their last appointment. We calculated cumulative incidence, incidence rate and fitted a multivariable Cox proportion hazards regression model to determine factors associated with LTFU.

**Results:** Of the 8,136 patients included in our sample, 3,606 (44.3%) started ART within seven days of HIV diagnosis. Females were 62.3%, median (interquartile range) age at start of ART was 30 (25, 37) years, 50.1% had access to a mobile phone, 54.0% had a baseline CD4 cell count of <350 cells/ml, 14.8% were in either WHO stage 3 or 4 at baseline and 75.9% had a normal body mass index (BMI). There were 1,207 cases of LTFU observed over 15953.0 person years at risk. The overall incidence rate (IR) of LTFU was 7.6 (95% CI=7.2-8.0) per 100 person years of observation (pyo). Cumulative incidence of LTFU increased with duration of follow up from 8.8% (95% CI=8.2-9.4%) and 12.0% (95% CI=11.2-12.7%) at 6 and 12 months, to 17.9% (95% CI=16.9-18.9%) and 20.1% (95% CI=18.9-21.3%) at 36, and 48 months respectively. Predictors of elevated risk of LTFU were; starting ART within 7 days of a positive diagnosis ((aHR) =1.39, 95% CI, 1.13-1.71), lack of access to a telephone set (aHR=1.60, 95% CI, 1.29-1.99) and baseline WHO clinical stage 3 or 4 (aHR =1.53, 95% CI, 1.11-2.11). Factors associated with a reduced risk of LTFU were; baseline age ≥25years, and having a BMI ≥ 30 (aHR =0.28, 95% CI, 0.15-0.51).

**Conclusion:** Initiation of ART within 7 days of an HIV diagnosis was associated with an elevated risk of loss to follow up. Steep ART initiation needs to be backed by enhanced adherence and retention counseling to reach the 2020 UNAIDS goals and beyond.

## Background and rationale

By the end of 2017, the World Health Organization (WHO) estimated that globally about 36.9 million people were living with HIV (PLHIV) and 1.8 million new infections occurred that year; over two thirds of the new infections were in Sub Saharan Africa (SSA) with about 50,000 in Uganda. During the same time, about 21.7 million (∼59%) of the PLHIV patients had been initiated on antiretroviral therapy (ART) [1]. To accelerate epidemic control, the United Nations Joint Program on HIV/AIDS (UNAIDS) set ambitious targets (90-90-90 campaign). One of the targets is to achieve 90% viral suppression in patients initiated on ART[2]. Whereas factors contributing to patients’ viral suppression are multi-factorial [3–8], retention of patients initiated on treatment and ensuring good adherence remain cornerstones to long term viral suppression and better treatment outcomes. The 2016 WHO and Ugandan Ministry of Health guidelines recommended start of ART regardless of CD4 and clinical stage of the disease[9,10]. The rationale is to arrest the disease before onset of opportunistic infections as well as accelerating achievement of the 2020 UNAIDS targets. Treatment as prevention studies demonstrated the effectiveness of ART in prevention of new HIV infections [11–14]. Therefore, scaling up ART coverage has public health benefits of reducing new infections through reduced community viral loads. Whereas achievement of the second and third UNAIDS targets demands timeliness in ART initiation, ensuring continuous engagement of patients with the health care system for periodic drug refills and running monitoring tests are critical to the success of this rapid ART scale-up. In spite of all this, loss to follow up of patients after ART initiation remains a great challenge. Systematic reviews of studies on the rapidly expanding ART programs in SSA illustrated that about 60-65% of patients were retained in HIV care at 2 to 3 years after starting ART [15,16]. In settings where patients start ART instantly after a positive HIV test, there is a possibility of offsetting the benefits associated with the immediate initiation when patients do not return to the HIV clinics. Patients need to perceive the clinical benefits of treatment continuation without interruptions. Previously, late presentation was common and HIV care and treatment guidelines stipulated that, only those presenting with WHO clinical stage III or IV of HIV/AIDS or had their CD4s decline to certain levels qualified for ART initiation. However, majority of patients currently diagnosed with HIV present in the early stages (WHO stage 1 or 2 or with CD4 cell count >500cell/ml) and initiate ART right away or shortly thereafter. Experiences from prevention of mother to child transmission (PMTCT) of HIV programs where instant ART initiation has widely been practiced indicate sub-optimal levels of ART adherence and retention of mothers in care[17, 18]. LTFU is associated with drug resistance, and comparatively poor long-term treatment outcomes, including mortality [19]. In resource constrained SSA countries, enhanced ART initiation will benefit from data characterizing retention of patients in typical clinical settings. To date however, a few studies have explored loss to follow up and associated factors in a typical HIV clinic practicing test and treat. Most of the data currently available are derived from implementation of test-and-treat in research settings. In this study, we set out to study the cumulative incidence and incidence rate of loss to follow up, and factors associated with loss to follow up in a primary healthcare clinic that has practiced test and treat since 2012.

## Methodology

### Study design

This was a retrospective cohort study utilizing data collected on patients who were diagnosed with HIV and enrolled into HIV care from January 2012 to December 2016 at Masaka regional referral hospital, -Uganda Cares’ clinic. A patient’s ART initiation date defined the beginning of follow up (time zero) and follow up period was from 01^st^ January 2012 to 31^st^ December 2016. Patients with documented transfer out status contributed follow up time up to the date of transfer out. Patients’ follow up ended if they died, transferred out to another HIV service delivery point, were LTFU or censored at 31^st^ December 2016.

### Study site and settings

Masaka regional referral hospital (MRRH), -Uganda Cares’ clinic serves as the main HIV outpatient department (OPD) clinic for the regional referral hospital. The total catchment population for MRRH currently exceeds 2,000,000 people (according to the Uganda population and housing census-UPHC, 2014), distributed in almost ten districts. The HIV clinic runs five days a week and by the end of 2016, more than 13,000 clients were active in care, with more than **86%** initiated on ART. The clinic serves patients of all characteristics including sex workers from the various hot spots of the Kampala-Masaka-Mbarara high-way and from neighboring fishing communities.

### HIV testing, linkage to care and initiation of ART in the study setting

At the beginning of 2012, provider initiated counseling and testing (PITC) was scaled up in MRRH. At the same time, voluntary counseling and testing (VCT) as well as home based HIV Counseling and Testing (HBHCT) outreaches were scaled up in the nearby villages. A serial testing algorithm was used during the study period. For screening; Determine™ HIV-1/2 (Alere Medical Company Limited, Chiba, Japan) and INSTI ® HIV-1/2 antibody test (Biolytical laboratories, Richmond, Canada). For confirmatory; Stat-Pak^®^ Dipstick (Chembio Diagnostic Systems, Medford, NY - USA) and Uni-Gold™ HIV (Trinity Biotech, Bray, Ireland) for tie breaking. Clients diagnosed with HIV within the hospital were enrolled into HIV care and encouraged to start on ART instantly or shortly after. During this period, a stand-alone desk (focal desk at the HIV clinic) was set up to fast track this. Patients diagnosed with HIV at the outreach sites were referred to this focal desk by trained counselors, to further aid and expedite ART initiation. Although the clinic begun piloting a T&T strategy at the beginning of 2012, it is important to note that this strategy wasn’t a true manifestation of T&T as illustrated by the treatment as prevention (TasP) group, but rather a process where the ART initiation process was expedited, with the preparatory counseling phase taking a maximum of one week. Under this T&T strategy, point of care CD4 cell count and TB assessment were done within a week to further determine ART eligibility. However, patients who declined ART instantly or within seven days were initiated on ART at a time of their convenience with a similar array of services as their counterparts who started ART instantly or shortly after. At the time and until June 2013, the ministry of health (MoH) policy to start ART was based on CD4 cell ≤350 cells/ml or WHO clinical stages 3 or 4 [20]. After this period, the guidelines changed to ART initiation based on CD4 cell count ≤500 cells/ml, WHO clinical stages 3 or 4 while “test and treat” was applicable to children, adolescents below 15 years and key populations [21].

### Study participants

We included all patients aged ≥18 years, tested and initiated on ART from 01^st^ January 2012 to 31^st^ December 2016 regardless of whether or not they tested at Masaka regional referral hospital. We excluded patients who transferred in from other HIV clinics because we were not able to confirm their HIV test and ART initiation dates with certainty, and patients with prior ART history (for example those that had ever used PEP since we could not ascertain the period when they were on medication). In addition, patients <18 years were excluded because they are children according to the Ugandan policy, but also because they do not decide on health service delivery options on their own but rather through their parents/care givers or guardians.

### Variables, data sources and measurement

The primary outcome was loss to follow up defined as failure of the client to show up at the Masaka clinic for at least 90 days from the date of their last scheduled appointment [37, 39] taking 31^st^ December 2016 as the reference date. We determined loss to follow up by comparing a patients’ most recent scheduled return visit date recorded into the electronic database with the reference date (31^st^ December 2016). The primary exposure (mode of treatment) was whether or not a patient was initiated on ART within seven days of HIV diagnosis. We defined test and treat (T&T) as patients who were initiated on ART within seven days of the HIV positive test; else, they were deferred. Other extraneous variables included; patients’ sex, age at ART initiation (determined by subtracting date of birth from the date of initiating ART), level of education, marital status, baseline CD4 cell count, baseline WHO stage, TB status at enrollment, ownership of a telephone set, and body mass index (BMI) calculated using the weight and height according to the formula BMI = weight/height(m)^2^. A study data extraction checklist with all study variables was piloted before data collection from the electronic database. The data were extracted in a Microsoft excel spread sheet, and cleaned. For incomplete and missing records, patients’ charts (source documents) were used to further clean the data set. A cleaned dataset was exported to Stata® version 13 for analysis.

### Statistical methods

We summarized patients’ characteristics by medians (interquartile range) for continuous variables and categorical characteristics were summarized by percentage. We limited determination of associations to only variables with complete data and only reported missing data at the descriptive stage. Comparison of continuous and categorical baseline characteristics was done by using t-tests and chi-square or Fisher exact tests. We analysed data at bivariate level to estimate crude estimates or predictors of associations and multivariable cox regression modelling to estimate adjusted predictors of time to loss to follow up. Multivariable model building involved a stepwise approach. We included variables with a p value of <0.2 at bivariate and dropped each turning out with a P>0.05 at multivariable model building. Insignificant variables at this stage but highlighted in previous literature as significantly associated with LTFU were included in the final model. The proportion hazards (PH) assumption was evaluated for each of the variable included in the final model. Kaplan Meier survival curves were used to determine patients LTFU. In all statistical tests, a 5% level of significance was assumed.

### Ethical considerations

This study was approved by the Makerere University School of Public Health Higher Degrees Research Committee. We also sought approval from the management of the HIV clinic, to allow us access to patients’ data. Program data routinely collected and entered into an electronic records management system (OpenMRS) was extracted without patients’ direct identifying information.

## Results

### Baseline characteristics

During the period January 2012 to December 2016, a total of 11,933 patients were extracted and 8,136 patients met the study inclusion criteria (Figure 1). Overall, there were more females (62.3%) than males. The median (IQR) age at start of ART was 30 (25-37), 20% started ART aged ≥40 years. About 86% had attained at least primary level of education. The proportion of those married and divorced was 41.0% and 20.9% respectively. A half (50.1%) had access to a telephone set and the mean (SD) weight at start of ART was 55 (10.7) Kgs. Overall median (IQR) CD4 cell count was 328 (180-490) cells/ml, with 301(171-440) cells/ml in the group starting ART ≤7 days and 352 (190-529) cells/ml in those who started after seven days. Our cohort had 14.8% of patients in either WHO clinical stage 3 or 4 at baseline, and 6.3% were suspected to have TB according to the initial TB assessment forms filled by clinicians. The proportion of patients who were started on a Tenofovir Disoproxil Fumarate (TDF) based regimen were 77.4% and just above three quarters (75.9%) had a normal baseline Body Mass Index (BMI) of 18.51-29.99.

**Figure 1:**
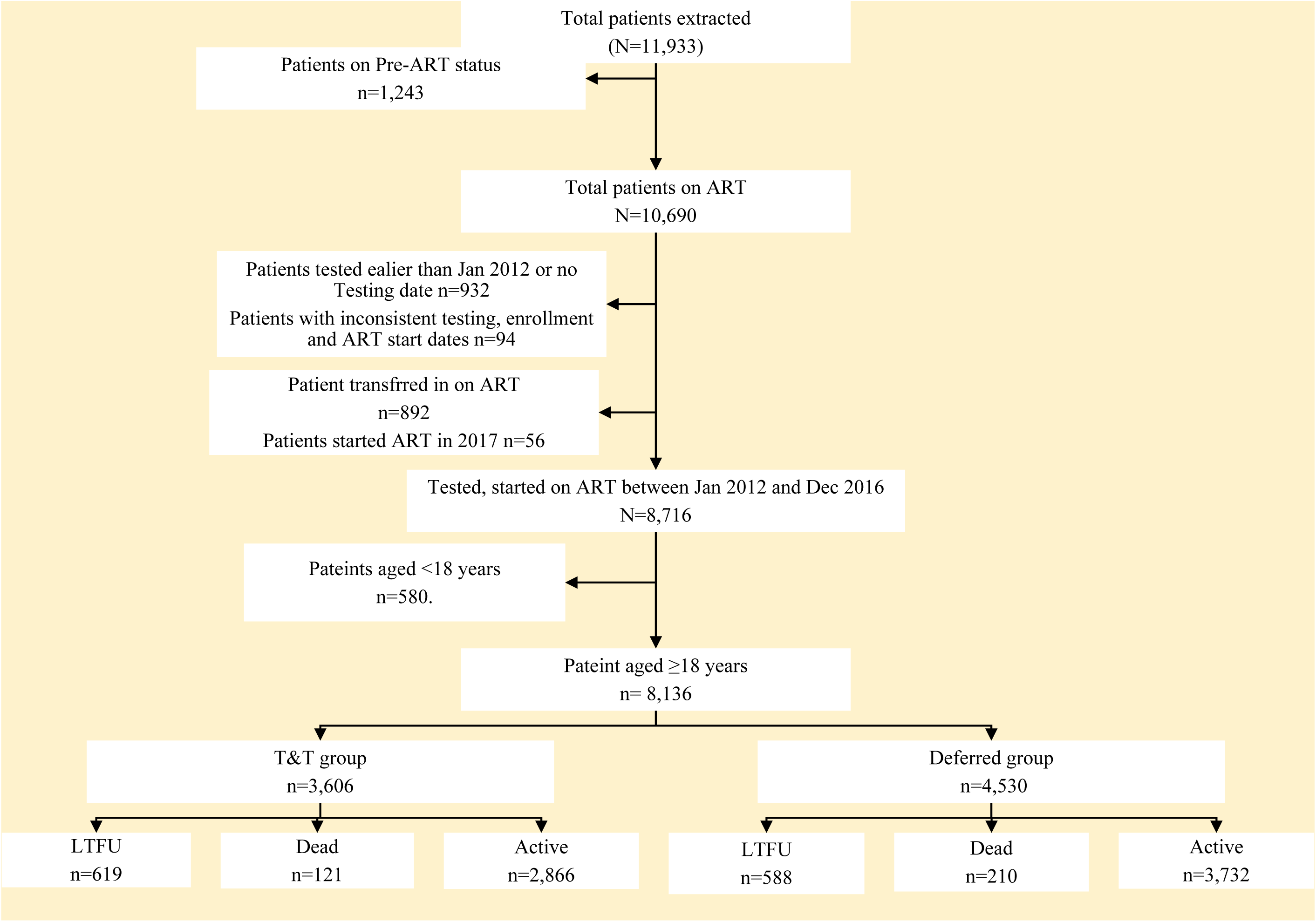
A flow diagram showing patients abstraction and inclusion into the study and their outcomes

### Study outcomes

#### a) Overall loss to follow up

There were 1,207 cases of LTFU observed over 15,953.0 person years at risk. The overall incidence rate (IR) of LTFU was 7.6 (95% CI=7.2-8.0) per 100 person years of observation (pyo). Cumulative incidence of LTFU at 6 months was 8.8% (95% CI=8.2-9.4%), 12 months 12.0% (95% CI=11.2-12.7%), 24 months 15.7% (95% CI=14.9%-16.6%), 36 months 17.9% (95% CI=16.9-18.9%) and 20.1% (95% CI=18.9-21.3%) at 48 months. Figures 2 and 3 illustrate the overall proportion of patients LTFU and cumulative incidence of LTFU by study group respectively. It can be observed that at all time points, the cumulative incidence of loss to follow up was significantly higher in the group that started ART within seven days of an HIV test compared to those who delayed.

**Figure 2:**
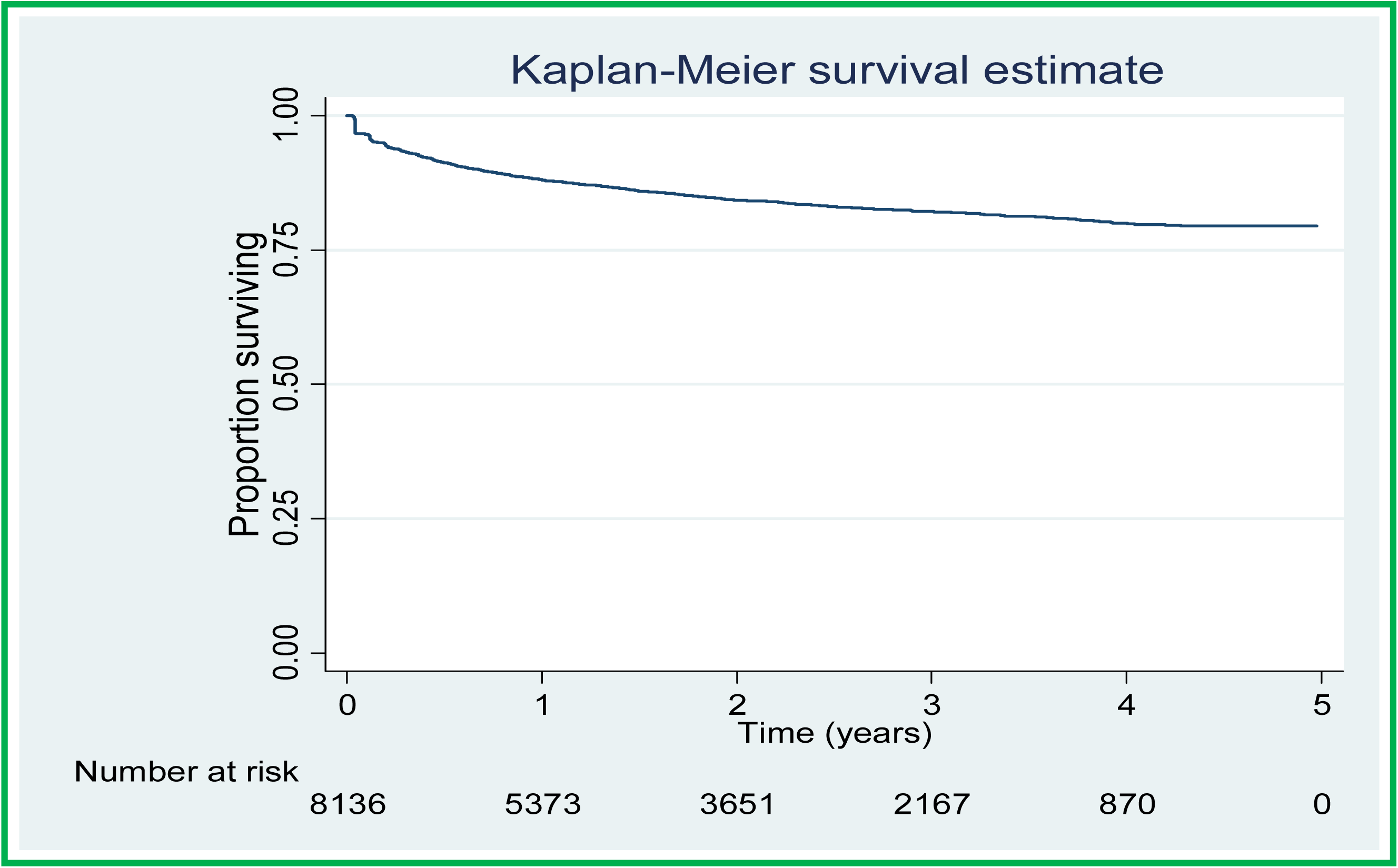
Overall Proportion of patients lost to follow up in the study period.

**Figure 3:**
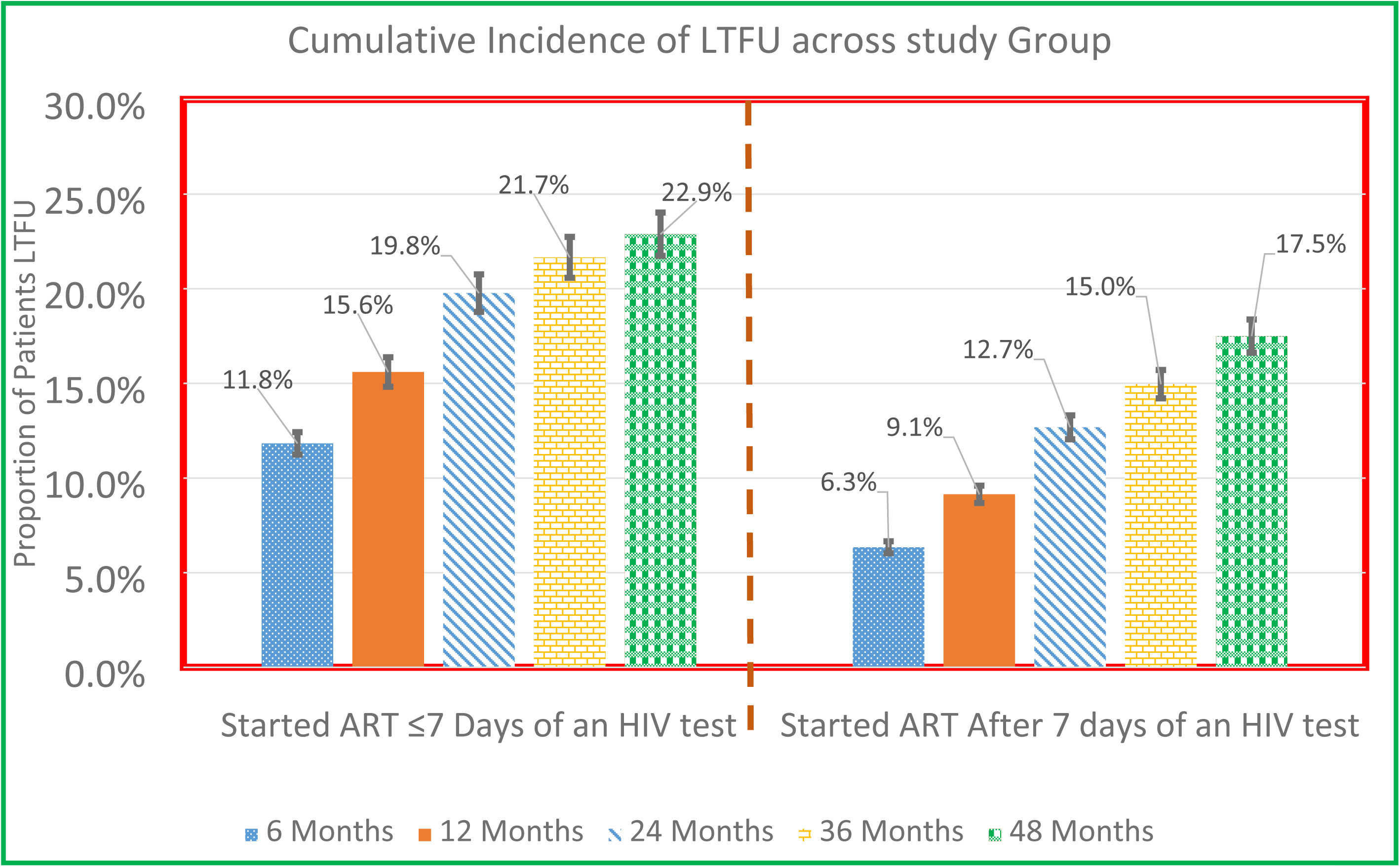
Cumulative Incidence of Loss to follow up by study group at different time points

#### b) Incidence rate of loss to follow up by patients characteristics

Table 2 indicates the IR of loss to follow up by patients’ characteristics. The incidence rate of LTFU was 10.9/100 pyo in the T&T group compared to 5.7/100 pyo in the delayed ART group. There was no difference in incidence rate of LTFU among males 7.4/100 pyo (95% CI, 6.7-8.1/100pyo) and females −7.7/100 pyo (95% CI, 7.2-8.3/100pyo).

**Table 1:** Patients background characteristics

| Characteristic (s) | Categories | Initiated on ART within 7Days |  | Initiated on ART after 7Days |  | All Groups |  |
| --- | --- | --- | --- | --- | --- | --- | --- |
|  |  | n | % | n | % | n | % |
| Sex | Male | 1242 | 34.4 | 1824 | 40.3 | 3066 | 37.7 |
|  | Female | 2364 | 65.6 | 2706 | 59.7 | 5070 | 62.3 |
| Age (years) | Median (IQR) | 30 (25-37) |  | 30 (25-37) |  | 30 (25-37) |  |
|  | 18-24 | 853 | 23.7 | 949 | 21.0 | 1802 | 22.1 |
|  | 25-29 | 945 | 26.2 | 1113 | 24.6 | 2061 | 25.3 |
|  | 30-39 | 1111 | 30.8 | 1576 | 34.8 | 2688 | 33.0 |
|  | 40-49 | 474 | 13.1 | 630 | 13.9 | 1105 | 13.6 |
|  | 50+ | 223 | 6.2 | 262 | 5.8 | 485 | 6.0 |
| Marital Status | Never Married | 248 | 6.9 | 351 | 7.8 | 599 | 7.4 |
|  | Married | 1586 | 44.0 | 1753 | 38.7 | 3339 | 41.0 |
|  | Divorced/Separated | 712 | 19.7 | 990 | 21.9 | 1702 | 20.9 |
|  | Widowed | 52 | 1.4 | 83 | 1.8 | 135 | 1.7 |
|  | Missing | 1008 | 28.0 | 1353 | 29.9 | 2361 | 29.0 |
| Education | None | 237 | 6.6 | 366 | 8.1 | 603 | 7.4 |
|  | Primary | 1827 | 50.7 | 2540 | 56.1 | 4367 | 53.7 |
|  | Secondary | 1094 | 30.3 | 1191 | 26.3 | 2285 | 28.1 |
|  | Post-secondary | 172 | 4.8 | 163 | 3.6 | 335 | 4.1 |
|  | Missing | 276 | 7.7 | 270 | 6.0 | 546 | 6.7 |
| Has Telephone | No | 1473 | 40.9 | 2588 | 57.1 | 4061 | 49.9 |
|  | Yes | 2133 | 59.2 | 1942 | 42.9 | 4075 | 50.1 |
| Baseline Weight (Kgs) | Mean (SD) | 56.6 (10.8) |  | 55.9 (10.6) |  | 55(10.7) |  |
|  | <45 | 337 | 9.5 | 473 | 10.5 | 810 | 10.0 |
|  | 45-60 | 2209 | 62.1 | 2831 | 62.7 | 5040 | 62.4 |
|  | 61+ | 1014 | 28.5 | 1211 | 26.8 | 2225 | 27.6 |
|  | Missing |  |  |  |  | 61 | 0.7 |
| Baseline CD4 cell count | Median (IQR) | 301 (171-440) |  | 352(190-529) |  | 328 (180-490) |  |
|  | <350 | 1949 | 59.4 | 2201 | 49.9 | 4150 | 54.0 |
|  | 350-500 | 793 | 24.2 | 944 | 21.4 | 1737 | 22.6 |
|  | >=500 | 538 | 16.4 | 1268 | 28.7 | 1806 | 23.5 |
| Baseline WHO clinical stage | 1&2 | 3150 | 87.7 | 3760 | 83.1 | 6910 | 85.2 |
|  | 3&4 | 441 | 12.3 | 763 | 16.9 | 1204 | 14.8 |
| TB status | No signs | 3283 | 91.0 | 3914 | 86.4 | 7197 | 88.5 |
|  | TB Suspect | 281 | 7.8 | 231 | 5.1 | 512 | 6.3 |
|  | TB Diagnosed | 5 | 0.1 | 274 | 6.1 | 279 | 3.4 |
|  | TB treatment | 19 | 0.5 | 100 | 2.2 | 119 | 1.5 |
|  | Missing | 18 | 0.5 | 11 | 0.2 | 29 | 0.4 |
| Baseline ART regimen | ABC based | 15 | 0.4 | 12 | 0.3 | 27 | 0.3 |
|  | AZT based | 480 | 13.3 | 1323 | 29.2 | 1803 | 22.2 |
|  | TDF based | 3108 | 86.2 | 3190 | 70.4 | 6298 | 77.4 |
|  | Other | 3 | 0.1 | 5 | 0.1 | 8 | 0.1 |
| Year of Starting ART | 2012 | 292 | 8.1 | 1056 | 23.3 | 1348 | 16.6 |
|  | 2013 | 538 | 14.9 | 1253 | 27.7 | 1791 | 22.0 |
|  | 2014 | 940 | 26.1 | 980 | 21.6 | 1920 | 23.6 |
|  | 2015 | 993 | 27.5 | 782 | 17.3 | 1775 | 21.8 |
|  | 2016 | 843 | 23.4 | 459 | 10.2 | 1302 | 16.0 |
| Baseline BMI | <18.50 | 407 | 21.0 | 639 | 21.4 | 1046 | 21.2 |
|  | 18.51-29.99 | 1480 | 76.3 | 2260 | 75.6 | 3740 | 75.9 |
|  | >=30 | 53 | 2.7 | 92 | 3.1 | 145 | 2.9 |

**Table 2:**
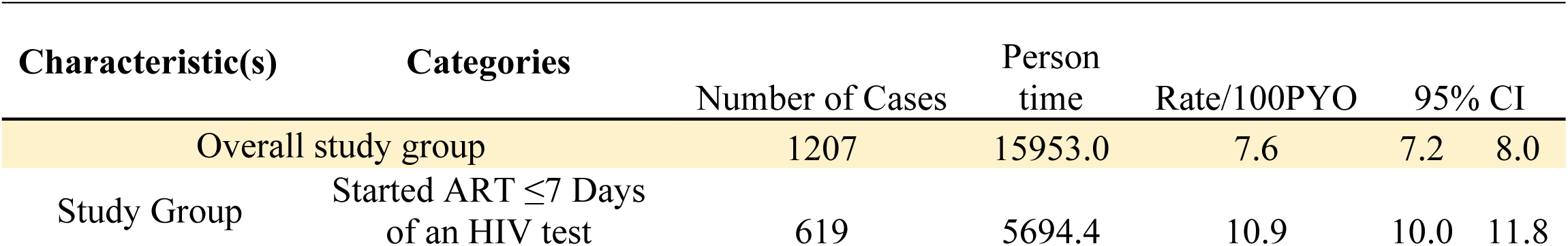
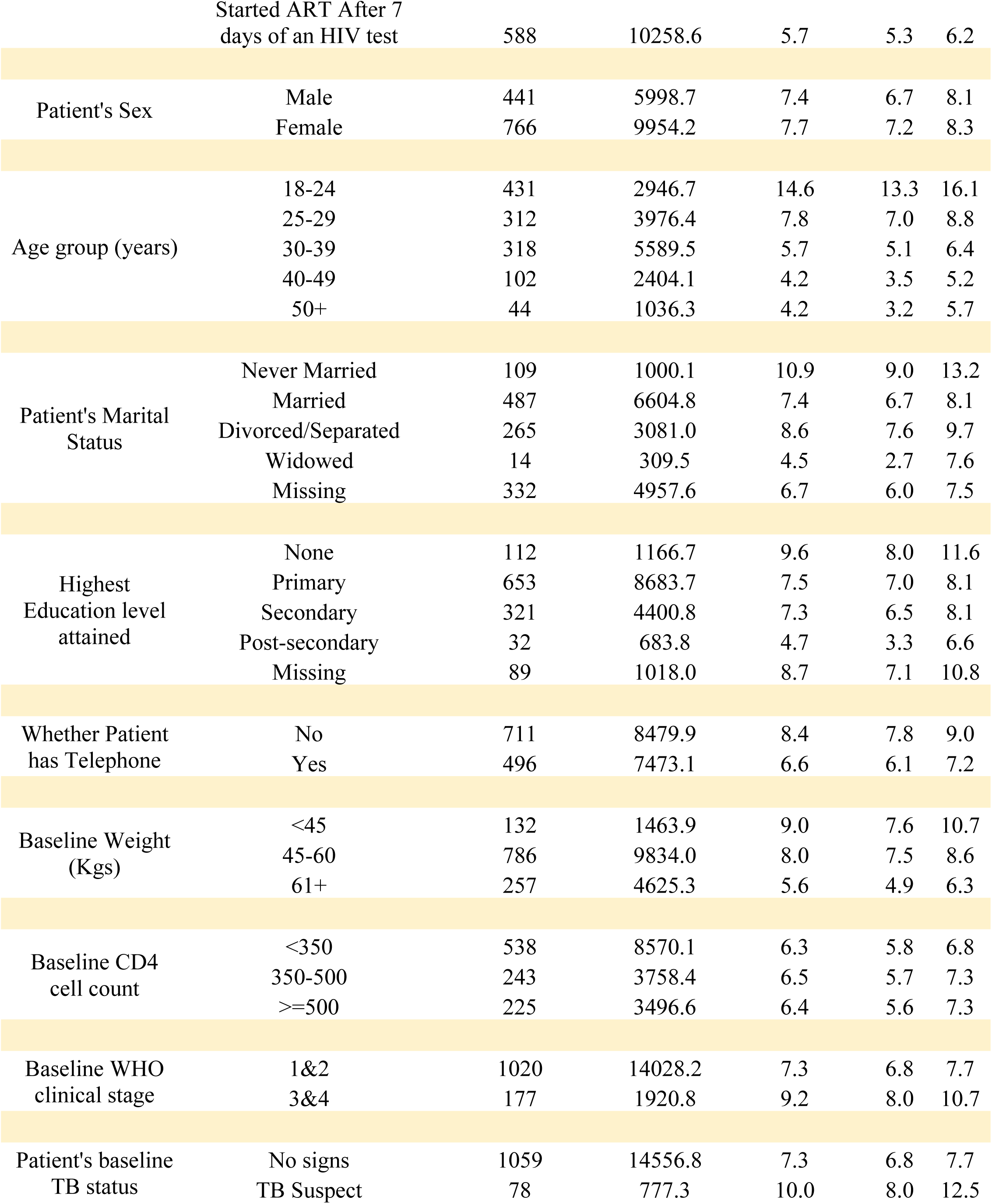
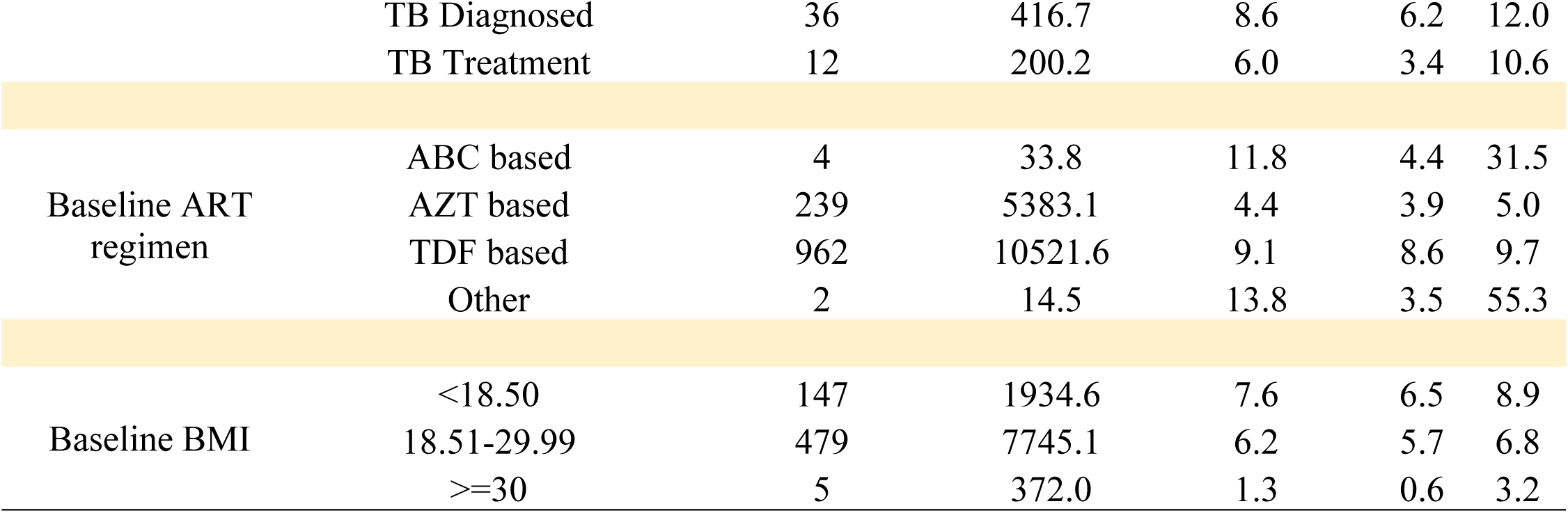
Incidence rate of Loss to follow up by Patients characteristics

The IR was highest at 14.6/100 pyo (95% CI, 13.3-16.1/100pyo) in patients that initiated ART in the age group 18-24 years and was lowest at 4.2/100 pyo (95% CI, 3.2-5.7/100pyo) among those aged ≥50 years. In patients with access to a telephone set, the IR was 6.6/100 pyo (95% CI, 6.1-7.2/100pyo) compared to 8.4/100 pyo (95% CI, 7.8-9.0/100pyo) in patients without access to a phone. Incidence rate of LTFU was 6.3/100 pyo (95% CI, 5.8-6.8/100pyo), 6.5/100pyo (95% CI, 5.7-7.3/100pyo) and 6.4/100 pyo (95% CI 5.6-7.3/100pyo) in patients with a baseline CD4 cell count of <350, 350-500 and ≥501 respectively. The incidence rate of LTFU was 7.3/100 pyo (95% CI, 6.8-7.7/100pyo) in patients who started ART with clinical disease classified as WHO stage 1 or 2 and this was statistically different from that of patients whose HIV disease was classified as either WHO stage 3 or 4 (IR=9.2/100pyo, 95% CI, 8.0-10.7/100pyo). Furthermore, IR was 7.6/100 pyo (95% CI, 6.5-8.9/100pyo), 6.2/100 pyo (95% CI, 5.7-6.8/100pyo) and 1.3/100 pyo (95% CI, 0.6-3.2/100pyo) in patients with a baseline body mass index of ≤18.50, 18.51-29.99 and ≥30 respectively.

#### c) Factors associated with loss to follow up

Table 3 depicts the unadjusted and adjusted hazard rates of factors associated with time to loss to follow up. None of the variables included in the final model violated the PH assumption. At bivariate, patients starting ART within seven days were 58% more likely to be lost to follow up (crude hazard rate (cHR)=1.58, 95% CI, 1.41-1.78).

**Table 3:** Factors associated with time to Loss to follow up.

| Characteristic (s) | Categories | crude<br>Hazard<br>Ratios | 95% CI | P<br>value | Adjusted<br>Hazard<br>Ratios | 95% CI | P value |
| --- | --- | --- | --- | --- | --- | --- | --- |
| Study Group | Started ART After<br>7 days of an HIV<br>test | 1.00 |  |  | 1.00 |  |  |
|  | Started ART ≤7<br>Days of an HIV<br>test | 1.58 | 1.41-1.78 | <0.001 | 1.39 | 1.13-1.71 | 0.002 |
| Patient's Sex | Male | 1.00 |  |  |  |  |  |
|  | Female | 1.05 | 0.94-1.18 | 0.390 |  |  |  |
| Age group (years) | 18-24 | 1.00 |  |  | 1.00 |  |  |
|  | 25-29 | 0.57 | 0.50-0.66 | <0.001 | 0.60 | 0.46-0.78 | <0.001 |
|  | 30-39 | 0.43 | 0.37-0.50 | <0.001 | 0.46 | 0.35-0.60 | <0.001 |
|  | 40-49 | 0.33 | 0.26-0.41 | <0.001 | 0.34 | 0.23-0.50 | <0.001 |
|  | 50+ | 0.32 | 0.24-0.44 | <0.001 | 0.23 | 0.12-0.42 | <0.001 |
| Patient's Marital<br>Status | Never Married | 1.00 |  |  | 1.00 |  |  |
|  | Married | 0.74 | 0.60-0.91 | 0.004 | 0.78 | 0.56-1.09 | 0.144 |
|  | Divorced/Separated | 0.83 | 0.66-1.03 | 0.092 | 1.03 | 0.73-1.44 | 0.879 |
|  | Widowed | 0.50 | 0.28-0.86 | 0.013 | 1.01 | 0.42-2.44 | 0.980 |
| Highest Education<br>level attained | None | 1.00 |  |  | 1.00 |  |  |
|  | Primary | 0.79 | 0.65-0.97 | 0.022 | 0.97 | 0.68-1.36 | 0.843 |
|  | Secondary | 0.76 | 0.61-0.94 | 0.010 | 0.72 | 0.49-1.05 | 0.085 |
|  | Post-Secondary | 0.49 | 0.33-0.73 | <0.001 | 0.47 | 0.22-1.02 | 0.056 |
| Whether Patient<br>has Telephone | Yes | 1.00 |  |  | 1.00 |  |  |
|  | No | 1.39 | 1.24-1.56 | <0.001 | 1.60 | 1.29-1.99 | <0.001 |
| Baseline CD4 cell<br>count | <350 | 1.00 |  |  | 1.00 |  |  |
|  | 350-500 | 1.03 | 0.88-1.20 | 0.710 | 1.06 | 0.83-1.36 | 0.619 |
|  | >=500 | 0.96 | 0.82-1.12 | 0.618 | 1.02 | 0.78-1.33 | 0.883 |
| Baseline WHO<br>clinical stage | 1&2 | 1.00 |  |  | 1.00 |  |  |
|  | 3&4 | 1.15 | 1.00-1.35 | 0.093 | 1.53 | 1.11-2.11 | 0.010 |
| Patient's baseline<br>TB status | No signs | 1.00 |  |  | 1.00 |  |  |
|  | TB Suspect | 1.21 | 0.96-1.52 | 0.103 | 1.12 | 0.77-1.64 | 0.551 |
|  | TB Diagnosed | 1.01 | 0.72-1.41 | 0.950 | 0.83 | 0.46-1.49 | 0.536 |
|  | TB Treatment | 0.76 | 0.43-1.35 | 0.351 | 0.37 | 0.12-1.21 | 0.101 |
| Baseline BMI | <18.50 | 1.00 |  |  | 1.00 |  |  |
|  | 18.51-29.99 | 0.85 | 0.71-1.03 | 0.092 | 0.88 | 0.69-1.13 | 0.323 |
|  | >=30 | 0.20 | 0.08-0.49 | <0.001 | 0.25 | 0.08-0.79 | 0.018 |

In the multivariable analysis, the risk of getting lost to follow up was 39% higher in patients that began treatment ≤7 days compared to those who began after seven days (adjusted Hazard ratios (aHR) =1.39, 95% CI, 1.13-1.71). Patients who started ART aged 18-24 years were more likely to get lost to follow up compared to all other age groups. In Comparison to patients aged 18-24 years at start of ART, those aged 25-29 years were 0.60 times (95% CI, 0.46-0.78) likely and patients aged ≥50 years were 0.23 times (95% CI, 0.12-0.42) likely. Compared to patients with no education, the risk of getting lost to follow was 0.97 times (95% CI, 0.68-1.36) in patients with primary level and 0.47 times (95% CI, 0.22-1.02) in patients whose baseline level of education was post-secondary, although statistical significance was borderline. Patients without access to a telephone set were 60% more likely to get lost compared to those with access to telephone (aHR =1.60, 95% CI, 1.29-1.99). Patients who started ART with HIV disease classified as WHO stage 3 or 4 were 53% more likely to get LTFU compared to those in WHO stage 1 or 2 (aHR =1.53, 95% CI, 1.11-2.11). Compared to patients with a baseline body mass index of <18.50, patients who were overweight at start of ART were 75% less likely to get LTFU (aHR =0.25, 95% CI, 0.08-0.79). There was no association between CD4 cell count at baseline, baseline TB assessment status and the risk of getting LTFU.

## Discussion

In this retrospective observation study of a primary healthcare clinic practicing test and treat, we observed a high cumulative incidence of loss to follow up. Four years after starting ART, one in every five patients that had started ART was lost to follow up. The proportion of LTFU was higher in patients that started ART within seven days of an HIV positive diagnosis than those who delayed ART. It is possible that patients had not yet received enough counselling and consequently not yet appreciated the benefit of starting ART when they were not yet “feeling sick”. Previously, patients were taken through a minimum of three counselling sessions, were required to bring a treatment supporter, and had to demonstrate understanding of long term treatment by answering questions after counselling [22]. We therefore speculate that probable lack of the perceived benefit to start ART on the same day of a positive diagnosis, coupled with inadequacies in the preparatory counseling could have led to the higher incidence of LTFU in the T&T group. In this cohort for example, of the 3606 patients that started ART within seven days, 52.1% initiated ART the same day of a positive HIV test. Addressing structural bottle necks including counseling after a positive HIV test and before ART initiation were identified as strategies for improving ART adherence and retention [23]. Inadequacies in pre-ART initiation counseling might compromise the patient’s perceived benefit for initiating ART instantly; similarly, during such a short while, key patients’ concerns that might affect long-term retention have not been exhaustively addressed.

The absolute differences in the proportions of patients LTFU in the subsequent time points and between treatment groups increased and peaked at 24 months. During 2012 and before, ART was not commonly provided in health centers at level III (Health Centers at level three). In the Ugandan setting, these provide outpatient services, maternity, general ward and laboratory. However, at the beginning of 2013 and going forward, most health centers at level III were ART accredited. This is suggestive that a large number of patients formerly at Masaka might have opted to receive ART services at these centers without formally seeking transfers/referrals. Self-transfers across ART programs have been illustrated to conceal the actual proportion of patients categorized as LTFU in a study in another similar setting[24]. In a similar realm, Masaka clinic serves as the main HIV OPD clinic for the regional referral hospital. We therefore speculate a possibility that patients diagnosed on wards and started ART instantly or within 7 days opted to receiving ART at health facilities nearest to their usual dwellings once they got better. Furthermore, these patients could have died after starting ART but were never reported given the passive nature of surveillance in our setting. Same day HIV diagnosis and ART initiation has widely been practiced under PMTCT (specifically under the Option B+). It has however been noted that retention in such settings has remained sub optimal. Moreover, initiation of ART on the same day of testing positive was independently associated with an elevated risk of loss to follow up in the initial months of starting ART [25–28]. A higher proportion of patients retained has been reported in a study in rural Uganda [29] and an almost comparable proportion in another study in Malawi [30] both at one year. Differences in proportions reported in these studies to ours could result out of methodological variations in determining loss to follow up as well as differences in the array of service delivery across the study populations. For example, Brown et al [30] evaluated retention in a streamlined care and universal test and treat model, a model designed to reduce patient barriers to care, unlike the typical and clinical setting in our study, while Jain et al [29] reported retention in asymptomatic patients with CD4 cell count restricted to ≥350 cells/ml..

Similar to another study in the same setting [31], retention rates in all other age groups were better compared to adolescence or being a young adult (18-24 years). Retention in adolescents and young adults should be an important subject given the rising rates of infection in this particular sub population [32] and high rates of viral un-suppression [33]. If not at school, adolescents and young adults are usually at conflict with work schedules and most times fail to make routine monthly schedules, a requirement in most ART clinics. Similarly, clumping ART services of adults together with those of adolescents and young adults might blur individualized adolescent and young adults’ needs. We anticipate that adolescents and young adults’ groups might benefit from differentiated care models that address individualized patients’ needs.

We observed that clients initiated on ART with WHO clinical stage categorized as either 3 or 4 were more likely to get LTFU compared to those staged 1 or 2. In asymptomatic patients initiated on ART in South Africa, the proportion of LTFU was low. Additionally, a reduced risk of death and improved retention rates were observed in a study assessing effectiveness of a streamlined model of care [29,30]. Patients categorized as being in stage 3 or 4 manifest with a higher likelihood of getting opportunistic infections, and so, will be bed ridden most of the time. This might make their continued engagement with the HIV clinic hard. Such patients are further at an increased risk of death especially in the first 6 months of ART due to severe immune reconstitution inflammatory syndrome and Cryptococcal meningitis [34,35]. It is therefore possible that they could have died shortly after starting ART and were never reported to the ART clinic.

We observed that patients with access to a telephone (mostly mobile) were less likely to get lost to follow up. In our setting, patients are sent short reminder text messages (SMS) before the clinic day and those who miss a clinic day are immediately called for a re-appointment. This cannot happen if one has no phone and so may lead to loss to follow up. Mobile phone technologies (mHealth), specifically SMS reminders have improved patient outcomes in other health service delivery settings [36–38] but were comparable to the standard of care for HIV retention in other settings [21,30]. It is however important to note that the level of interaction between provider and patient, and subset of activities under mHealth greatly determine the effectiveness of particular interventions. Therefore, interactive SMS reminders alone might not improve patient outcomes when compared to a combined strategy of SMS reminders, home visits and direct phone calls to patients.

We did not find a statistically significant relationship between CD4 cell count and LTFU. This finding was also observed by Jain et al [26] but contrasts results of studies in other similar settings [40–42]. One possibility for the contrast could be differences in the determination of LTFU as well as the differing CD4 cell count thresholds used across the studies. A patient was classified LTFU if they had spent at least 180 days without picking their ART from the HIV clinic after a scheduled visit [41]. Berheto et al and Honge et al classified patients as LTFU if they failed to pick their ART after 90 days from the last scheduled date [40,42], a similar definition to that used in our study. In the papers written by Berheto et al and Honge et al, an elevated risk of LTFU was noticeable in patients that had a baseline CD4 cell count of <200 cells/ml [37,39] while a similar risk was observed in patients with a CD4 cell count of >200 cells/ml by Mberi et al [41]. In comparison to our cohort, we categorized CD4 cell count as <350 cells/ml, 350-500 cells/ml and >500 cells/ml. Nevertheless, changes in WHO and in-country treatment guidelines over the course of study period might have resulted into almost similar immunological patients starting ART in the T&T and deferred groups.

### Limitations

Our study had limitations that should be taken into account while interpreting these findings. First, we utilized already collected data used for routine patient management. Such data presents with lots of gaps and sometimes may not present the rigor to warrant scientific research. Whereas this particular clinic is not a research site, it is part of the east African International epidemiology database to evaluate AIDS (IeDEA) consortium. As such, there are inherent data validation rules within the database and daily data cleaning to guarantee a certain degree of data correction and collation. Secondly, as it is in most HIV programs, there is a passive nature of surveillance and follow up of patients. There is therefore a possibility of having determined and regarded patients as lost to follow up in Masaka when they are actually in HIV care and receiving treatment somewhere else. This therefore, might have resulted into over estimation of the cumulative and incidence rates of lost to follow up in our study. There is however a dedicated team at the facility that does contact tracing/case navigation for clients who miss clinic appointments, and we think this might minimize on this misclassification. Lastly, the nature of data collected was limited. Some of the many health system (human resource, waiting time, distance) and socio-economic factors (type of work, socio contacts, HIV disclosure) known to affect loss to follow up were not studied. The effect of such under the current study settings remained unknown. We anticipate that, examining the effect of these under a test and treat setting could better inform ART programing and policies towards boosting retention.

## Conclusions

Our study shows that initiation of ART within 7 days of a positive HIV test is associated with an elevated risk of loss to follow up in the long run. Steep ART initiation needs to be backed by enhanced intensive adherence and retention counseling for improved long term patient outcomes by 2020 and beyond. Our findings further categorize the risk of getting lost to follow up in patients sub groups. This is beneficial to HIV service providers to recognize patients that require enhanced support in this era of test and treat.

## Acknowledgements

We extend our sincere gratitude to Mathew Ssemakadde (Data manager at Masaka Uganda Cares clinic) for his tireless efforts getting us the data from OpenMRS. We are indebted to Dr Cecilia Nattembo, for her insightful reviews and corrections. We also wish to thank the management of Masaka Uganda cares clinic for allowing us use their routine program data without hesitation. We are especially grateful to all the patients from whom big volumes of data are collected routinely.

## Authors’ contributions

### Conceived and designed the experiments

Julius Kiwanuka, Noah Kiwanuka. **Performed the experiments:** Julius Kiwanuka, Noah Kiwanuka, Jacinta Mukulu Waila, and Jonathan Kitonsa. **Analysed the data:** Julius Kiwanuka. **Wrote and approved the manuscript:** Julius Kiwanuka, Jonathan Kitonsa, Jacinta Mukulu Waila, Methuselah Kahungu Muhindo, and Noah Kiwanuka.

